# Sequestration and functional diversification of cyanogenic glucosides in the life cycle of *Heliconius melpomene*

**DOI:** 10.1101/723973

**Authors:** Érika C. P. de Castro, Rojan Demirtas, Anna Orteu, Carl Erik Olsen, Mohammed Saddik Motawie, Márcio Z. Cardoso, Mika Zagrobelny, Søren Bak

**Author notes:** Corresponding author: Søren Bak^a^.

## Abstract

*Heliconius* butterflies are highly specialized in *Passiflora*, laying eggs and feeding as larvae only on these plants. Interestingly, *Heliconius* butterflies and *Passiflora* plants both contain cyanogenic glucosides (CNglcs). While feeding on specific *Passiflora* species, *Heliconius* m*elpomene* larvae are able to sequester simple cyclopentenyl CNglcs, the most common CNglcs in this plant genus. Yet, aromatic, aliphatic, and modified CNglcs have been reported in *Passiflora* species and they were never tested for sequestration by heliconiine larvae. As other cyanogenic lepidopterans, *H. melpomene* also biosynthesize the aliphatic CNglcs linamarin and lotaustralin, and their toxicity does not rely exclusively on sequestration. Although the genes encoding the enzymes in the CNglc biosynthesis have not yet been fully biochemically characterized in butterflies, the cytochromes P450 CYP405A4, CYP405A5, CYP405A6 and CYP332A1 are hypothesized to be involved in this pathway in *H. melpomene*. In this study, we determine how the CNglc composition and expression of the putative P450s involved in the biosynthesis of these compounds vary at different development stages of *Heliconius* butterflies. We also established which kind of CNglcs *H. melpomene* larvae can sequestered from *Passiflora*. By analysing the chemical composition of the haemolymph from larvae fed with different *Passiflora* diets, we observed that *H. melpomene* is able to sequestered prunasin, an aromatic CNglcs, from *P. platyloba*. They were also able to sequester amygdalin, gynocardin, [C^13^/C^14^]linamarin and [C^13^/C^14^]lotaustralin painted on the plant leaves. The CNglc tetraphyllin B-sulphate from *P. caerulea* was not detected in the larval haemolymph, suggesting that such modified CNglcs cannot be sequestered by *Heliconius*. Although pupae and virgin adults contain dihydrogynocardin resulting from larval sequestration, this compound was metabolized during adulthood, and not used as nuptial gift or transferred to the offspring. Thus, we speculate that dihydrogynocardin was catabolized to recycle nitrogen and glucose, and/or to produce fitness signals during courtship and calling. Mature adults had a higher concentration of CNglcs than any other developmental stages due to intense *de novo* biosynthesis of linamarin and lotaustralin. All *CYP405As* were expressed in adults, whereas larvae mostly expressed *CYP405A4*. Our results shed light on the importance of CNglcs in Heliconius biology and for their coevolution with *Passiflora.*

## 1. INTRODUCTION

Toxicity is undoubtedly an important feature in the evolution of *Heliconius* butterflies, whose bright and colourful wings advertise their unpalatability to possible predators. (Sheppard et al., 1985; Reed et al., 2011). Cyanogenic glucosides (CNglcs) are the most abundant toxic compounds in *Heliconius* butterflies and also in *Passiflora* plants, their coevolutionary partners. *Heliconius* exclusively lay eggs and feed as larvae on *Passiflora* plants which are normally avoided by other insect herbivores due to their CNglc content (Ehrlich and Raven, 1964; Engler-Chaouat and Gilbert, 2007; de Castro et al., 2018).

*Heliconius* butterflies biosynthesize the aliphatic CNglcs linamarin and lotaustralin through a pathway hypothesized to be orthologous between butterflies and *Zygaena* moths (Chauhan et al., 2013). In the burnet moth *Z. filipendulae*, this pathway is encoded by the genes *CYP405A2, CYP332A3* and *UGT33A1* (Jensen et al., 2011). The two cytochromes P450 (P450s), CYP405A2 and CYP332A3, convert the amino acids valine and isoleucine into their respective cyanohydrins, which are stabilized with a glucose residue added by the UDP-glycosyltransferase (UGT) UGT33A1, forming linamarin or lotaustralin. In *H. melpomene*, the precursors and intermediates of the CNglc pathway have been confirmed to be the same as in *Z. filipendulae* (Davis and Nahrstedt, 1987) and accordingly the P450s catalysing the reactions are thought to be orthologues to the moth pathway: *CYP405A4, CYP405A5, CYP405A6* and *CYP332A1* (Chauhan et al., 2013) (Figure 1). In the *Z. filipendulae* a single CYP405A catalyses the first reaction, however, in *H. melpomene* there are three closely related CYP405A that putatively may catalyse the reaction The three *HmCYP405As* are 80-88% identical at the coding sequence and tandemly clustered in the same genome scaffold (Zagrobelny et al., 2018). An orthologues UGT could not be identified from the *H. melpomene* genome, perhaps due to different UGTs being recruited into the pathway in butterflies and moths (Zagrobelny et al., 2018).

**Figure 1.**
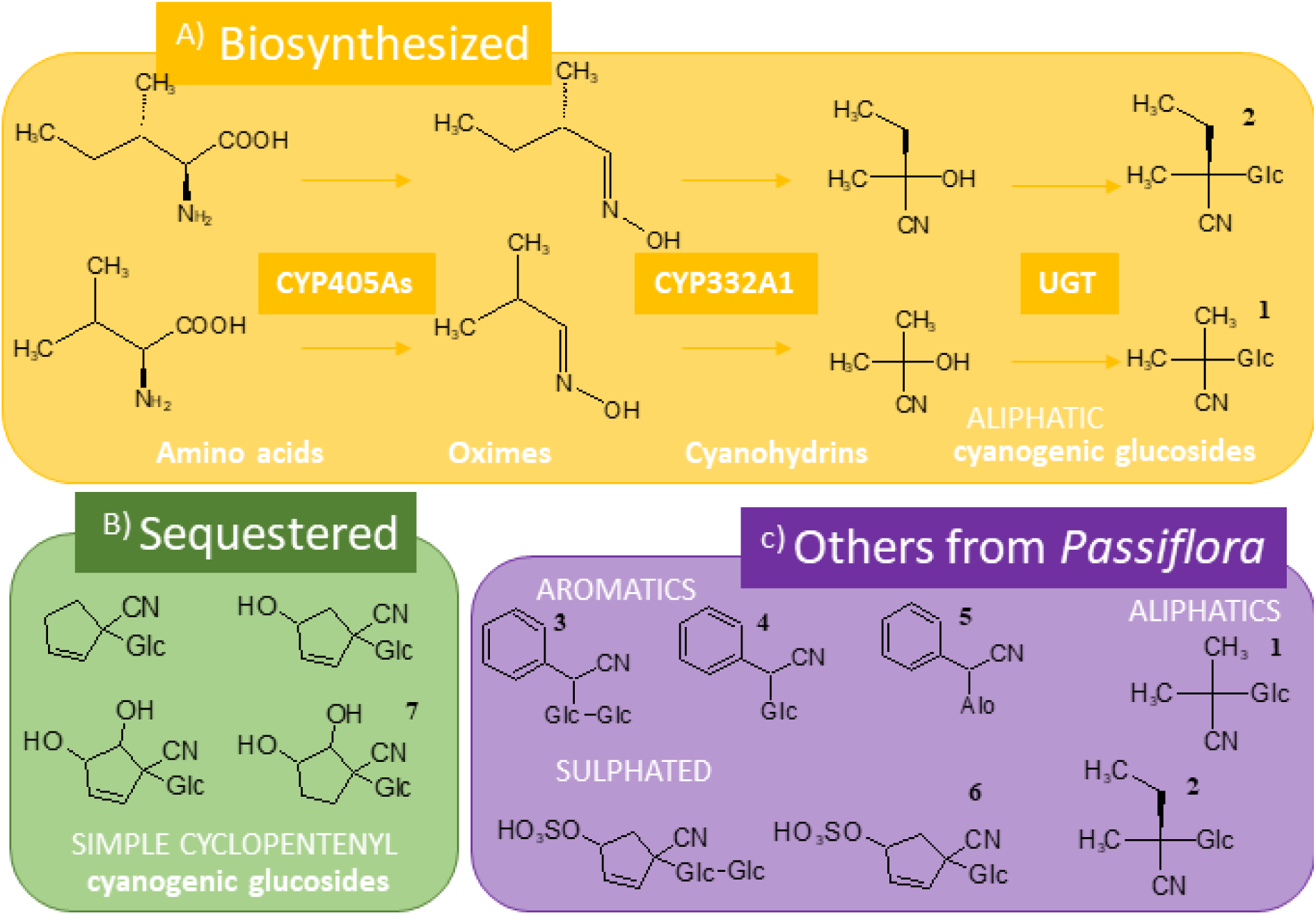
Cyanogenic glucosides (CNglcs) in *Heliconius melpomene* and *Passiflora plants.* A) Biosynthesized CNglcs: *H. melpomene* biosynthesize the aliphatic CNglcs linamarin^1^ and lotaustralin^2^; P450s hypothesized to be involved in this pathway are CYP405A4, CYP405A5, CYP405A6 and CYP332A1. B) CNglcs previously known to be sequestered: As in other heliconiines, larvae can also sequester cyclopentenyl cyanogenic glucosides from *Passiflora*, such as dihydrogynocardin^7^. C) Others CNglcs found in the genus *Passiflora:* Many other CNglcs, such as amygdalin^3^, prunasin^4^, passiedulin^5^ and tetraphyllin B-sulphate ^6^, have been found in *Passiflora* hosts utilized by *H. melpomene* and it is not known if they are sequestered.

Plants from the *Passiflora* genus also produce CNglcs, and it has previously been demonstrated that CNglcs biosynthesized from cyclopentenyl glycine by *Passiflora* can be sequestered by *Heliconius* (Engler et al., 2000; Engler-Chaouat and Gilbert, 2007). Some of these cyclopentenyl CNglcs, most likely remnants from larval feeding, were even found in adults of several *Heliconius* and heliconiine species, suggesting that these compounds are kept after pupation (de Castro et al. 2019) (Figure 1). However, an enormous diversity of CNglc structures is present in the *Passiflora* genus, which raise the question whether *Heliconius* larvae could sequester other CNglcs apart from the ones with a cyclopentenoid structure.

More than 60 different kinds of CNglcs have been found in over 2,500 plant species (Møller, 2010) – of which 27 were reported in the *Passiflora* genus alone, including some structures that seems to be produced only by *Passiflora* (Jaroszewski et al., 2002). This indicates that *Passiflora* is an evolutionary hot-spot for evolution of novel CNglc structures. CNglcs derived from the non-protein amino acid cyclopentenyl glycine are the most common CNglcs in *Passiflora*, they are only found in Passifloraceae and five other closely related plant families. In some *Passiflora* species these cyclopentenyl CNglcs are modified with additional unusual sugars and sulphate groups, such as passibiflorin and tetraphyllin B, respectively (Figure 1). The aliphatic CNglcs linamarin and lotaustralin are present in some *Passiflora* species (Spencer et al., 1986), whereas few others contain the aromatic CNglcs prunasin, amygdalin, and passiedulin, of which has only been reported in *P. edulis* (Christensen and Jaroszewski, 2001; de Castro et al., 2019).

Spencer (1988) hypothesized that *Passiflora* plants diversified their CNglc profile in response to the herbivory of *Heliconius* and closely related genera. By producing CNglcs that cannot be sequestered, a *Passiflora* species would decrease its value as a host for these larvae. However, to confirm this hypothesis it is necessary to establish which CNglc structures *Heliconius* larvae can actually sequester from their hosts. Additionally, *Heliconius* must be able to avoid the degradation of these compounds during sequestration, since plant tissue disruption caused by herbivory will put CNglcs in content with β-glucosideses, which will lead to their conversion into volatile breakdown products (cyanide and aldehydes/ketones)(Pentzold et al., 2014a).

Since *Heliconius* butterflies both synthesize and sequester large amounts of CNglcs, it is clear that these metabolites are important to their biology. Yet, the roles of these compounds in the life history of these butterflies have not been fully elucidated. For instance, the defensive function of CNglcs is well established in plants and it has been hypothesized to confer protection to *Heliconius* as well, but so far only circumstantial evidence has been presented (Schappert and Shore, 1999; Gilbert, 1991; Cardoso, 2019). Other roles for CNglcs in lepidopterans have been suggested by Cardoso and Gilbert, (2007), who based on detected of cyanide emission from the spermatophores of several *Heliconius* species, found evidence for that these butterflies also use CNglcs as nuptial gifts, and by Zagrobelny et al., (2007b), who suggested that CNglcs are used as nitrogen and sugar storage molecules by *Zygaena* moths. Nevertheless, the dynamics of CNglcs in the life history of *Heliconius* butterflies has not been fully explored.

In this study, we investigated in detail, not only how CNglcs content fluctuates during the whole life cycle of a *Heliconius* species, but also elucidated it from a standpoint of sequestration versus biosynthesis. We examined which CNglcs *H. melpomene* can sequester by exposing larvae to different CNglc-containing diets and analysed the compounds sequestered into their haemolymph. We also established how cyanogenesis varies during the life-cycle of *H. melpomene*, by characterizing the CNglc profile from pupae to mature adults and in the development of their offspring. Furthermore, we examined the expression of the putative enzymes involved in the biosynthetic pathway at different life stages. Finally, spermatophores and testes were also analysed to establish which CNglcs *H. melpomene* can use as nuptial gifts

## 2. Methods

### 2.1. Plants samples

Leaf discs (1 cm^2^) of *P. caerulea, P. edulis* and *P. platyloba* were collected in tubes containing 500 mL methanol 80% (V/V) + formic acid 0.1% (V/V) and boiled for five min. Samples were collected from six month old plants that had not been exposed to oviposition or herbivory. The sampled plants were afterwards used in the feeding experiments.

### 2.2. Butterflies and breeding conditions

The butterflies were hatched from pupae bought from Costa Rica Entomological Supplies. Pupae were placed in small cages (0.4m × 0.3 m × 0.15 m) and kept under controlled conditions (24-28 °C, 80% RH, 14h day/10h night). Cages were examined every day, and newly eclosed butterflies were transferred to a larger cage for reproduction or sampled as freshly emerged imagines. The cage for reproduction (2 m × 1.5 m × 1m) was supplied with flowering *Lantana* plants for adult feeding (nectar and pollen) and young *Passiflora* plants for oviposition. Adult feeding was supplemented with plastic flowers containing 20% honey solution (v/v) with 1% (m/v) fresh willow pollen (Percie Du Sert, UK). For the life cycle experiment, only plants of *P. edulis* were used for oviposition and freshly laid eggs were sampled. First instar larvae (1-3) were collected one week from the date when the first eggs were laid. Late instar larvae (4-5) were identified and collected when the skin of their dorsum was completely white (Brown, 1981). Mature adults were collected when larvae of their offspring were reaching the final instars (∼4 weeks after eclosion). All collected samples were frozen in liquid nitrogen and stored at – 80 °C. Specimens destined for isolation of spermatophore and testes were collected in plastic bags and kept in the freezer (– 20 °C) until dissection.

### 2.3. No-choice larval feeding experiments

Plants from different *Passiflora* species were located in the cage for reproduction with freshly emerged *H. melpomene* imagines. The *Passiflora* plants utilized were from the species *P. caerulae, P. edulis* and *P. platyloba*, which have previously been used to breed *H. melpomene* under insectary conditions in our facilities. The cage had four plants of each species. The sulphated cyclopentenyl CNglc Tetraphyllin B sulfate is present in *P. caerulae*, whereas *P. edulis and P. platyloba* contain the aromatic CNglcs prunasin and amygdalin. Eggs were left to hatch on the plants where they were laid and raised until the fourth instar. Larvae were then isolated in individual plastic boxes containing a moistened paper tissue and a leaf of their previous host-plant species. The plastic boxes were cleaned and supplied with fresh leaves every day.

To confirm and compare the larvae’s ability to sequester cyclopentenyl and aliphatic CNglcs, some larvae raised on *P. platyloba* were fed with leaves painted with 5 mg of gynocardin or with 3 mg of labelled [^14^C/^13^C] linamarin and lotaustralin. Gynocardin was previously synthesized by Jaroszewski et al., (2002) and the radiolabelled [^14^C/^13^C] linamarin and lotaustralin was produced in our lab as described by Zagrobelny et al., (2014). *P. platyloba* was used because it is the favourite host plant of *H. melpomene* in our facilities, and because it lacks aliphatic and cyclopentenyl CNglcs. Painted leaves were left to dry for at least 2 h, before being added to feeding boxes. Leaves of *P. caerulea*, which do not contain amygdalin, were painted with 10 mg of this compound and used to feed some of the larvae reared on this species. After 24h, the frass of each larva was collected and larvae were prepared for haemolymph sampling (see section 2.4). Frass pieces from a larva reared on *P. edulis* were separated in saline buffer and photographed under an EZ4 (Leica) stereo microscope. A minimum of five larvae fed on each type of diet.

### 2.4. Dissection of larvae and imagines

Larvae were starved for 4 hours, rinsed twice with distilled water, and subsequently kept for 30 seconds in a chamber with dry-ice to be anesthetized prior to dissection. A foreleg from each larva was removed with chirurgic scissors and the haemolymph collected with a pipette trough this incision. The haemolymph samples were weighed in microtubes and frozen in liquid nitrogen. Gut and skin samples were also collected. Five gut samples (not frozen) were used for pH measurements of the gut lumen.

Imagines were kept in the freezer for a minimum of 72 h before spermatophores and testes were collected. Abdomens were separated and immersed in saline buffer under an EZ4 (Leica) stereo microscope. Testes were collected from males and spermatophores retrieved from the female bursa. Samples were weighed in microtubes and frozen in liquid nitrogen.

### 2.5. Liquid chromatography–mass spectrometry (LC-MS/MS)

Extractions, LC-MS conditions and analyses were executed as described in Castro et al., 2019

### 2.6. Gene-expression analyses

Total RNA was extracted from frozen eggs, larvae, pupae, and adults of *H. melpomene* utilizing the RNeasy Mini Kit (Qiagen), including digestion with the RNase-Free DNase Set (Qiagen). cDNA was synthesized using the iScript cDNA Synthesis Kit (Bio-Rad) using 500 ng of total RNA as template. qRT-PCR reactions were conducted with DyNAmo Flash SYBR Green qRT-PCR Kit (Thermo Fisher Scientific) with 150 ng of cDNA and specific primers (Table S1). No-template added controls were also included. The reactions were run in a CFX384 Touch™ qPCR system (Bio-Rad) following the conditions described in Table S2, and the data were analysed in the native software. The method ΔΔCt was used to calculate the relative expression of the genes of interest: *CYP405A4, CYP405A5, CYP405A6* and *CYP332A1.* In this method, the expression of the genes of interest were normalized against the housekeeping gene Elongation Factor 1 alpha (EF1a), and all the samples standardized using the values found in the egg samples as a reference.

To validate the qRT-PCR results, gene expression analyses were also performed using published RNA sequencing (RNA-seq) data of *H. melpomene* larvae (gut) from Yu et al. (2016) and adults (abdomen) from Walters et al. (2015). Given that the data came from two separate projects, each dataset was analysed independently following the same methodology. First, low quality base and adapter trimming, and quality control of the reads were performed using TrimGalore! (Krueger, 2015). Reads were then mapped to the *H. melpomene* genome (H.mel2.5)(Davey et al., 2016) using STAR (version 2.5.0a) (Dobin et al., 2013). Read counts were produced using featureCounts (Liao et al., 2013) and then normalised by library size (TMM normalisation) and gene length (log2 RPKM) using the edgeR package in R (Robinson et al., 2010). Differences in gene expression between genes of the same dataset were assessed with pairwise *t*-tests with Bonferroni correction for multiple testing.

### 2.7. Statistical analyses

To compare the total concentration of CNglcs between larvae fed different diets, and between samples from different life stages, we used one-way ANOVA with post-hoc Tukey tests. Two-way ANOVA was employed for comparison of P450 genes putatively involved in the CNglc biosynthesis during the life cycle of *H. melpomene*. When appropriate, post-hoc Tukey tests were employed. All analyses were performed in R.

## 3. Results

### 3.1. Variation in the CNglc profile of larvae on different diets

To investigate how larval diet impacts the CNglc profile of *H. melpomene* and which CNglcs they are able to sequester from *Passiflora*, larvae were raised to fourth instar on *P. caerulea, P. edulis* or P. *platyloba*. The chemical profile of these plants were analysed *a priori* of the experiment: *P. caerulea* produces tetraphyllin-B sulphate, *P. edulis* prunasin, and *P. platyloba* prunasin and amygdalin (Figure 2a). The total CNglc concentration of *P. platyloba* is higher than the others (Tukey, p=0.0002 against *P. caerulea*; p=0.00019 against *P. edulis*). None of these plants produce linamarin and lotaustralin.

**Figure 2.**
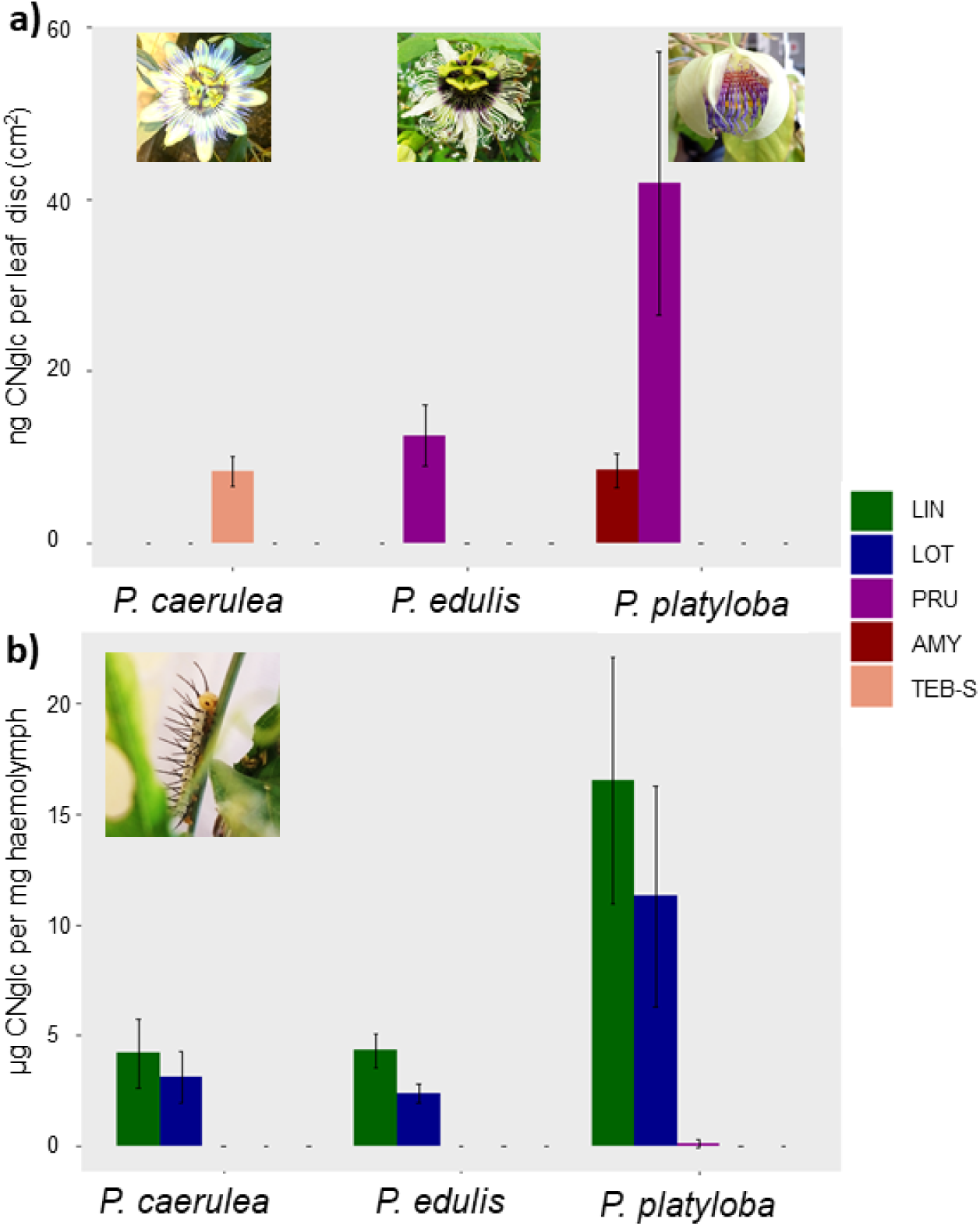
**a)** Concentration of CNglcs (ng cm^2^ leaf disc) in the *Passiflora* species used in this study. The plants contained tetraphyllin-B-sulfate (TEB-S), prunasin (PRU) and amygdalin (AMY**); b)** CNglc concentration in the hemolymph of *H. melpomene* larvae raised on the *Passiflora* plants in previous panel. Larvae biosynthesize linamarin (LIN) and lotaustralin (LOT) and sequester small amount of PRU from *P. platyloba*. Error bars correspond to standard deviation (N=4-5)

Haemolymph of larvae raised on the three different diets were collected and analysed by LC-MS/MS to establish the larval CNglc profile. CNglc accumulation in the haemolymph of *H. melpomene* differed between the three *Passiflora* diets (ANOVA, Df=2 F= 14.01, p= 0.0035) (Figure 2b). Larvae raised on *P. platyloba* had a higher total concentration of CNglcs than *P. edulis* (Tukey, p=0.0065) and *P. caerulea* (Tukey, p=0.0053), the latter two did not differ from each other (Tukey, p=0.9907). All larvae contained linamarin and lotaustralin, which does not originate from sequestration but from biosynthesis. Larvae reared on *P. platyloba* biosynthesized more linamarin and lotaustralin than on other plant diets, which is probably associated with the nutritional value of this species as a host. Sequestration of CNglcs was only observed in larvae fed on *P. platyloba*, which were only able to sequester small amounts of the aromatic CNglc prunasin into their haemolymph, but not its corresponding diglucoside amygdalin. Sequestration of tetraphyllin B-sulphate from *P. caerulea*, and prunasin from *P. edulis* was not detected.

In order to increase the repertoire of compounds ingested by the larvae, leaves of *P. platyloba* painted with either gynocardin or [^14^C/^13^C] radiolabelled linamarin and lotaustralin, and *P. caerulea* painted with amygdalin were used (Table 1). The radiolabelled linamarin and lotaustralin was used to enable discrimination between the amounts sequestered and biosynthesized. When *P. caerulea* leaves were painted with amygdalin (10 mg per leaf) and ingested by larvae, a small amount of amygdalin was sequestered into their haemolymph (0.12 µg/mg) (Table 1), but most of it was excreted in the frass (3.06 µg/mg) (data not shown). In contrast, larvae intensively sequestered gynocardin into their haemolymph when fed on *P. platyloba* painted with this cyclopentenyl CNglc (5 mg per leaf). Larvae fed on *P. platyloba* leaves painted with an even ratio of radiolabelled linamarin and lotaustralin (5 mg per leaf) had a higher total concentration of CNglcs due to intense sequestration of these compounds. Interestingly, we observed that radiolabelled linamarin was sequestered to a much higher extent than radiolabelled lotaustralin (69.7 ± 10.4 µg /mg linamarin versus 2.4 ± 0.2 µg /mg lotaustralin), even though they were supplied in equal proportion. Larvae on this diet synthesized more lotaustralin than linamarin (28.14 ± 9.9 versus 19.4 4± 6.1 µg/mg, respectively), most likely in an attempt to keep the relative level of linamarin to lotaustralin in balance. Sequestration of prunasin was also increased in comparison to *P. platyloba* without additional compounds (2.74 ± 2.94 and 0.39 ± 0.27 µg/mg, respectively), thought this difference was mainly caused by an elevated concentration of prunasin in one particular larva. This is the first time to our knowledge that the sequestration of both aromatic and aliphatic CNglcs are reported in *Heliconius.*

**Table 1.** Concentration of different CNglcs (µg/mg) in the haemolymph of *H. melpomene* larvae fed on *Passiflora* leaves painted with different CNglcs.

|  | Present in the non-painted diet |  |  | Present **only** in the painted diet |  |  |  |
| --- | --- | --- | --- | --- | --- | --- | --- |
| | LIN | LOT | PRU | AMY | GYN | [$^{14}\text{C}/^{13}\text{C}$] LIN | [$^{14}\text{C}/^{13}\text{C}$] LOT |
| *P. caerulea* + AMY | 2.37<br>$\pm 1.24$ | 1.3<br>$\pm 0.73$ | N/A | 0.12<br>$\pm 0.05$ | N/A | N/A | N/A |
| *P. platyloba* + GYN | 11.14<br>$\pm 4.72$ | 7.86<br>$\pm 3.31$ | 0.39<br>$\pm 0.27$ | 0 ± 0 | 8.05<br>$\pm 6.20$ | N/A | N/A |
| *P. platyloba* + [$^{14}\text{C}/^{13}\text{C}$]LIN+LOT | 19.44<br>$\pm 6.14$ | 28.14<br>$\pm 9.96$ | 2.74<br>$\pm 2.94$ | 0 ± 0 | N/A | 69.7<br>$\pm 10.46$ | 2.42<br>$\pm 0.24$ |

Note: See Figure 2b, for CNglc concentration in the haemolymph of larvae raised on non-painted *P. caerulea* and *P. platyloba*.

### 3.2. Adaptations to sequester CNglcs

Frass from *H. melpomene* larvae were composed of small leaf pieces (Figure S1), indicating that they perform leaf-snipping while eating. This feeding mode help keeping CNglcs intact during feeding, as it avoids that ingested plant β-glucosidases comes into contact with CNglc to degrade them. The average gut lumen pH of *H. melpomene* larvae was 9.73 ± 0.02. This alkaline gut probably inhibits the activity of ingested plant β-glucosidases and further protects the CNglcs from degradation, as was also shown in the *Zygaena* moths (Pentzold et al., 2014a)

**Figure S1.**
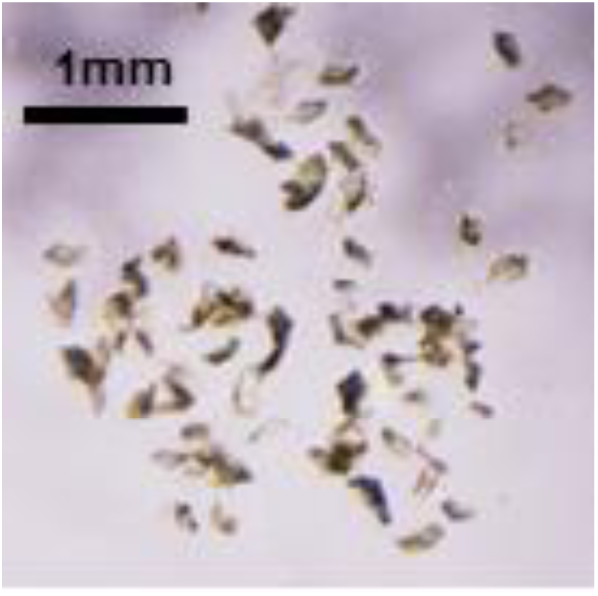
Light microscopy pictures from the frass of *Heliconius melpomene* larvae.

### 3.3. CNglcs in the life-cycle of H. melpomene

The amount and type of CNglcs were examined in different life stages of *H. melpomene* to establish how this varies during the life cycle. Total concentration of CNglcs varied significantly between different developmental stages (ANOVA, Df=5, F= 16.58, p= 9.74e-08) (Figure 3a). Pupae and virgin adults (newly emerged) had a similar total concentration of CNglcs (Tukey, p=0.865). The amount of dihydrogynocardin, sequestered during larval feeding, was in the same range as the biosynthesized linamarin and lotaustralin. The highest CNglc concentration was observed in mature adults which had 10 times more linamarin and lotaustralin relative to dihydrogynocardin. This reduction in the dihydrogynocardin content from virgin to mature adults indicates that the sequestered dihydrogynocardin is consumed during adulthood. The increase in linamarin and lotaustralin in mature adults in comparison to virgins reflects the continuous biosynthesis at this stage. Only linamarin and lotaustralin were present in the eggs, suggesting that these two CNglcs, but not dihydrogynocardin, could be transferred to the offspring. Eggs had a similar overall CNglc concentration to parental pupae (Tukey, p=0.953) and virgin adults (Tukey, p=0.429). As the progeny were raised on *P. edulis*, it was not possible to sequester CNglcs during larval feeding and larvae had to rely exclusive on biosynthesis to acquire their cyanogenic defences (Figure 2a). Early instar larvae had the lowest CNglc concentration of all the life stages, whereas late instar larvae had similar total concentration of CNglcs to parental pupae (Tukey, p=0.1461828) showing biosynthesis of linamarin and lotaustralin was later intensified during larval development of *H. melpomene*. Eggs and larvae did not contain dihydrogynocardin, demonstrating that dihydrogynocardin is neither transferred from parents to progeny nor biosynthesized (Fig 3a+b). Linamarin and lotaustralin, but not dihydrogynocardin, were found in both testes and the spermatophore of *H. melpomene* butterflies. This corroborates earlier results showing that they Heliconius use CNglcs as nuptial gifts (Cardoso and Gilbert, 2007) (Figure 4b).

**Figure 3:**
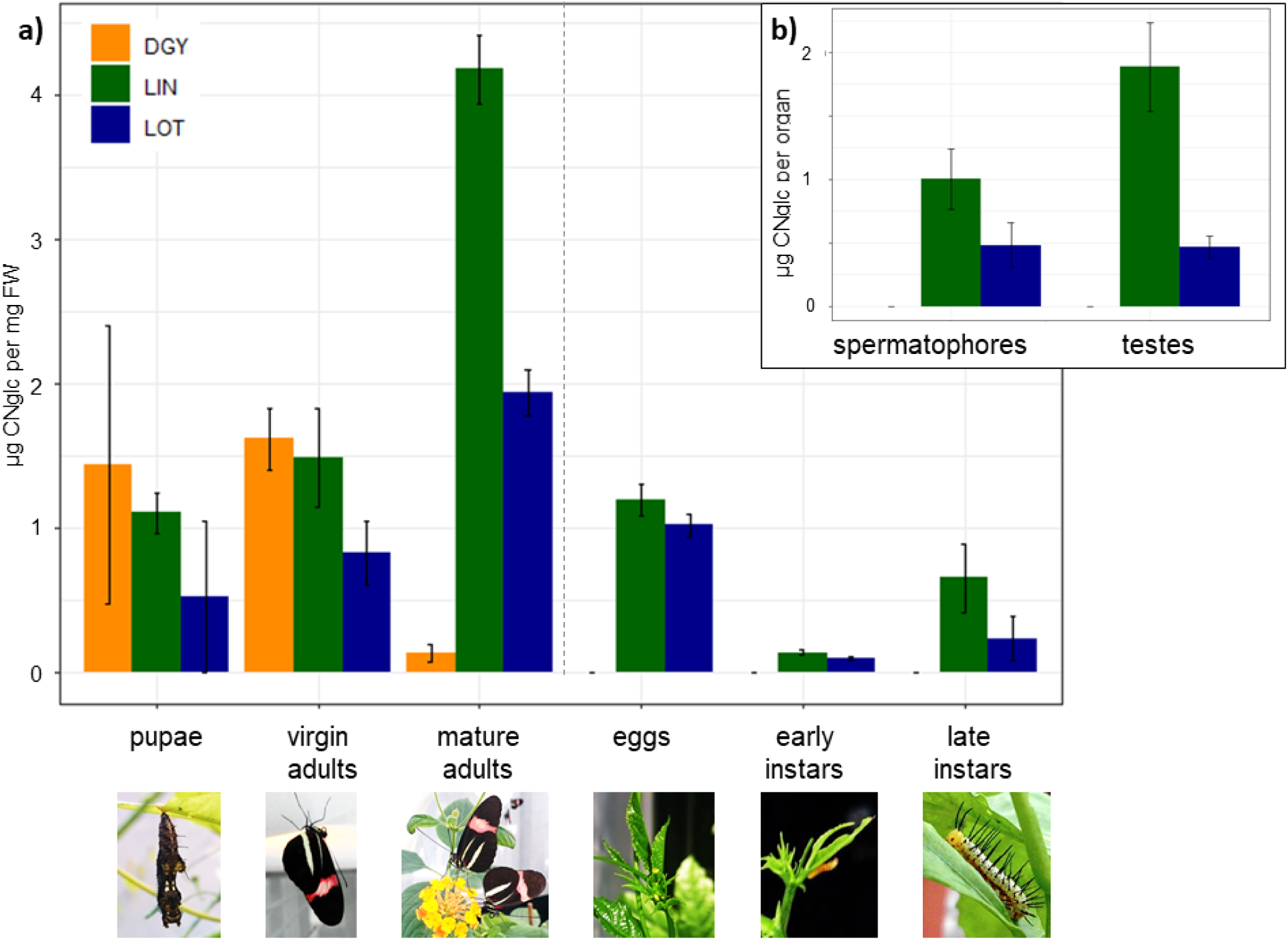
**a)** CNglc concentration in pupae, virgin and old adults of *H. melpomene* and their progeny (eggs, early and late instar larvae); **b)** CNglc content of spermatophores and testes. Error bars correspond to standard error (N=5-10). Note: Nuptial gifts refer to spermatophore only, not to testes.

**Figure 4.**
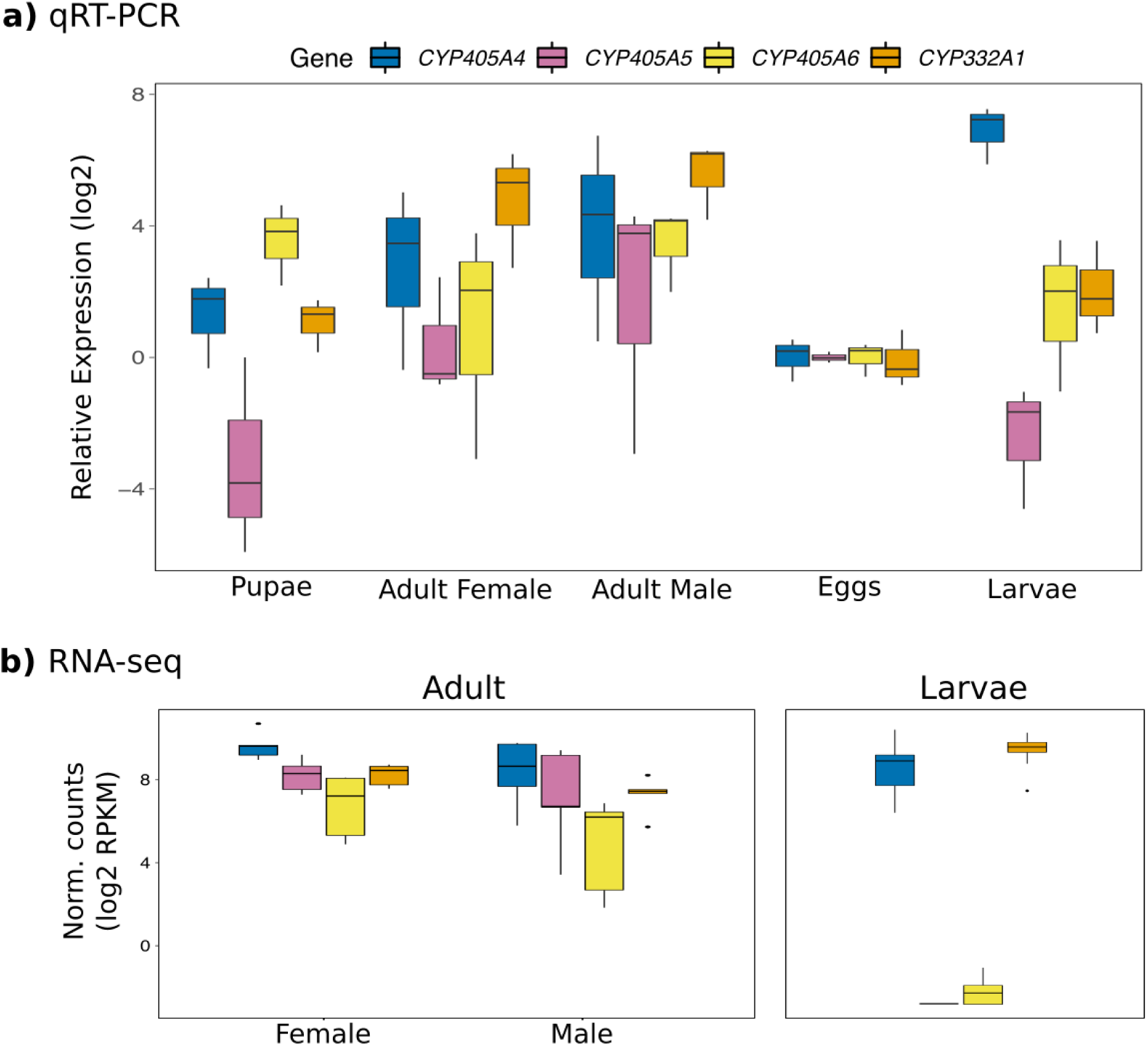
Gene expression of P450 genes putatively involved in the CNglcs biosynthesis during the life cycle of *H. melpomene*, calculated from **a)** qRT-PCR and **b)** RNA-seq samples. Relative expressions in qRT-PCR were calculated relative to the expression level in eggs using the 2^−ΔΔCt^ method with EF1a as a housekeeping gene. RNA-seq results are expressed as log2 RPKM normalised counts.

### 3.4. Gene expression and CNglc biosynthesis in the life cycle of H. melpomene

The expression of the P450s putatively involved in the CNglc biosynthesis was measured in eggs, larvae, pupae and freshly emerged adults of *H. melpomene* by qPCR (Figure 4a). Generally, lepidopteran eggs acquired their chemical defences from their parents and do not biosynthesize them (Eisner et al., 2000; Fürstenberg-hägg et al., 2014; Mason and Singer, 2015). In agreement with this, our qPCR results show the lowest Cts for the analysed genes at the egg stage. Hence, the gene expression in the other life stages were normalized relatively to the expression levels in eggs.

Across all life stages and candidate genes the highest expression was observed for *CYP405A4* in larvae (P < 0.01, pairwise t-test with Benjamini & Hochberg correction). In this life stage, *CYP405A4* was on average expressed more than 2.2 fold higher than any of the other three candidate genes (*P* < 0.01). In general, adults had higher expression levels of the analysed P450 genes than pupae – especially of *CYP405A4* and *CYP332A1* – although there were no significant differences in the gene expressions among adult males or females. In all the life stages, *CYP405A5* always had the relative lowest expression.

To corroborate our qRT-PCR results, gene expression analyses were performed using publicly available RNA sequencing (RNA-seq) samples from *H. melpomene* 5^th^ instar larvae (gut) (Yu et al., 2016) and adult males and females (abdomen; Figure 4b)(Walters et al., 2015). Both the qRT-PCR and RNA-seq data indicate that out of the three *CYP405As, CYP405A4* is preferentially expressed in larvae, while *CYP405A5* and *CYP405A6* are not. Similar to our qRT-PCR results, *CYP405A4* had higher expression than *CYP405A5* and *CYP405A6* (*P* < 1e-25, *t*-test, Bonferroni correction). However, in this case, higher expression of *CYP332A1* was also detected (*P* < 1e-26), which could be due to the difference in age of the larvae used in the analysis: 1^st^ instar larvae in the qRT-PCR and 5^th^ instar larvae in the RNA-seq. As observed in the qRT-PCR results, all genes were expressed in both females and males, and no significant differences could be detected among them in each sex, with the exception of *CYP405A4*, which was more highly expressed then *CYP405A6* in adult females.

## 4. Discussion

### 4.1. The CNglc profile in Heliconius varies according to the CNglc profile of their Passiflora hosts

Although plants produce defence compounds to keep herbivores a bay, specialized herbivores can overcome these defences and even utilize them for their own benefit, selectively taking them up into their own tissues through the process of sequestration.

In this study, it was observed that larvae of *H. melpomene* can sequester not only cyclopentenyl CNglcs, but also aromatic and aliphatic CNglcs (Figure 2b and Table 1). Sequestration of sulphated cyclopentenyl CNglcs (tetraphyllin B-sulphate) from *P. caerulea* was not detected (Figure 2b), which suggested that addition of a sulphate group to CNglcs in *Passiflora* could be a recent modification to avoid sequestration by heliconiines, as previously hypothesized (de Castro et al., 2019). Larvae did not sequester prunasin from *P. edulis* and amygdalin from *P. platyloba* (Figure 2b), even though we demonstrated that they have the machinery to sequester these compounds when they are painted on leaves in high concentration. This could indicate that sequestration of aromatic CNglcs by most *Heliconius* larvae is only possible when they are highly concentrated in *Passiflora* leaves, as e.g. prunasin in *P. platyloba* (Figure 2b and Table 1).

When larvae were fed with *P. platyloba* leaves painted with radiolabelled linamarin and lotaustralin, *H. melpomene* larvae sequestered over 10 times fold more radiolabelled linamarin than lotaustralin, despite the even ratio presented to the larvae. To compensate for this difference, we speculate that they synthesized more lotaustralin than linamarin to uphold a certain ratio (Table 1). Similarly, *Z. filipendulae* larvae tend to maintain a linamarin:lotaustralin ∼70:30 ratio in their haemolymph even when raised on diets containing more lotaustralin than linamarin (Zagrobelny et al., 2007a). In plants and also in insects, linamarin and lotaustralin are synthesized by similar enzyme complexes and the relative affinity of the enzymes for the respective substrate amino acids, valine and isoleucine, and their availability defines which of them will be produced (Zagrobelny et al., 2008). Therefore, the first P450 of the biosynthetic pathway, CYP79 for plants and CYP405 for Lepidoptera, is important for the maintenance of the linamarin:lotaustralin ratio (Jensen et al., 2011; Fürstenberg-Hägg et al., 2014). Our results suggest that the stoichiometry of the reaction catalysed by the CYP405 in *H. melpomene* is regulated by the concentration of the final products of the pathway, linamarin and lotaustralin, similar to in *Zygaena* moths (Zagrobelny et al., 2008; Jensen et al., 2011).

Larvae of *H. melpomene* are very inefficient at sequestering aromatic CNglcs from *Passiflora* plants: only trace amounts of prunasin were sequestered from *P. platyloba* while uptake of aromatic CNglcs from *P. edulis*, which produced ∼3 fold less, was not detected at all (Figure 2a and 2b). Small amounts of amygdalin were only sequestered when it was artificially supplied in large amounts (10 mg per leaf) painted on *P. caerulea* leaves (Table 1). This result is contrary to what is observed in *Z. filipendulae* which can sequester large amounts of both linamarin, lotaustralin and prunasin, even though only the first two are naturally occurring in its food plant (Pentzold et al., 2015b). The low efficiency of sequestration of aromatic CNglcs in *Heliconius* could be caused by the absence of a transporter that can take up aromatic CNglcs. Since aromatic CNglcs are not very common within the *Passiflora* genus (reported only in three species), there would be no selection in favour of a sequestration apparatus for aromatic CNglcs in heliconiines.

*H. melpomene* produced more linamarin and lotaustralin when fed *P. platyloba* leaves than when reared on *P. caerulea* or *P.* edulis (Figure 2b). Curiously, *P. platyloba* and *P. edulis* have both aromatic CNglcs (Figure 2a). This suggests that not only the CNglc profile, but also e.g. the nutritional value of the host plant affects cyanogenesis in *Heliconius.* Leaves of *P. platyloba* seem to have a thin cuticle, they are soft and with light green coloration, even in mature leaves (personal observations), which is the opposite of *P. edulis* leaves that are glossy dark green on the adaxial side of the leaf (Wosch et al., 2015). Consequently, *P. platyloba* could have a higher nutritional value, facilitating more CNglc biosynthesis in *H. melpomene.* The main aim of this study was to analyse the impact of the *Passiflora* CNglc profile on the CNglc profile in *Heliconius*. Accordingly, measurements related to larval development, such as growth rate, developmental time, weight gain and food consumption, were not evaluated in this experiment, but would be very useful to include in future experiments based on our results.

### 4.2. Heliconius larvae avoid degradation of CNglcs during feeding

To avert self-poisoning, plants often glycosylate their defence compounds (benzoxazinoids, cyanogenic and iridoid glucosides, glucosinolates, etc.) and store them separately from β-glucosidases (Pentzold et al., 2014b). During herbivore or pathogen attack, tissue disruption will result in these two components coming into contact and the removal of the sugar residue will elicit the toxicity. Alkaline midgut pH is very common in Lepidoptera (Berenbaum, 1980), and since plant β-glucosidases are normally inhibited under this condition, it allows Lepidoptera to tolerate many toxic glycosylated compounds. *H. melpomene* larvae perform leaf-snipping during feeding (Figure S1) and have an alkaline gut lumen pH. These two features have also been observed in *Z. filipendulae* and are considered adaptations to circumvent the degradation of CNglcs by plant β-glucosidases (Pentzold et al., 2014a). By biting off leaf pieces without macerating them, larvae minimize the release of compartmentalized β-glucosidases, and consequently CNglc hydrolysis and production of toxic cyanide. Any residual β-glucosidase activity left on the leaf edges or in the leaf pieces will be inhibited by the alkaline midgut pH. Larvae of the fall webworm (*Hyphantria cunea*) also has an alkaline midgut environment which allow them to feed on black cherries without trigging their cyanogenic potential (Fitzgerald, 2008).

Amelot et al., (2006) demonstrated that larvae of *Heliconius erato* released less hydrogen cyanide than *Spodoptera furgiperda* when feeding on *Passiflora capsularis* (5.8 fold difference). *S. furgiperda* is a leaf-chewer (Nabity et al., 2012), not a snipper as *Heliconius*, and its midgut environment is also less alkaline (pH= 8.5-9.5)(Alfonso et al., 1997) Thus, the minor cyanide release during *H. erato* feeding is likely due to the two adaptations reported here.

### 4.3. Mature adults of H. melpomene are highly cyanogenic

In the life-cycle of *H. melpomene*, mature adults have a higher concentration of CNglcs than pupae and newly emerged butterflies due to their increased content of linamarin and lotaustralin (Figure 3a). This indicates that biosynthesis of linamarin and lotaustralin is intensified during the adult stage, and corroborates earlier findings that butterflies of *H. melpomene* are less cyanogenic at eclosion than after 7 to 28 days (Nahrstedt and Davis 1985). Intensification of CNglcs biosynthesis in adults might be a result of all three *CYP405A* being expressed at this stage (Figure 4).

Contrary to *H. melpomene, Z. filipendulae* have higher concentrations of CNglcs as final instar larvae than as eggs or adults (Zagrobelny et al., 2007b). As larvae, *Z. filipendulae* biosynthesize linamarin and lotaustralin, and also sequester the same compounds from their host *Lotus corniculatus. Z. filipendulae* stops the expression of the genes involved in the CNglc biosynthesis as they start to pupate and they are only re-activated in females close to eclosion. Accordingly, male adults only have the CNglcs that originates from accumulation during the larval period (Fürstenberg-Hägg et al., 2014). Turn-over of CNglcs was also observed during pupation, contributing to the lower concentration of these compound in adult *Zygaena* moths in comparison with larvae (Zagrobelny et al., 2007a)

Most aposematic butterflies sequester defence compounds as larvae and utilize this reservoir in their own defence as adults (Nishida, 2002). *H. melpomene* not only synthesize defence compounds as pupae and adults, but also intensifies the process during adulthood. This could be a reflection of a much longer adult life-span in *Heliconius* species compared to other butterflies, including other heliconiines, and is perhaps associated with their pollen-feeding behaviour (Gilbert, 1972). For example, *H. charithonia* butterflies can survive more than 100 days, while *Dryas iulia*, another heliconiine that does not supplement its nectar-diet with pollen, has an adult life-span of maximum 35 days (Boggs, 1981). *Heliconius* are more unpalatable than other heliconiine butterflies and thus it has been hypothesized, that they have more CNglcs due to their pollen-feeding behaviour, which among other things provide them with the amino acids necessary for the biosynthesis of these compounds. Nevertheless, recently emerged *Heliconius* butterflies have overall similar total concentration of CNglcs to other heliconiines (de Castro et al., 2019), which suggests that *Heliconius* might become more toxic than other heliconiines as they become mature adults (3-4 weeks after eclosion). Corroborating this hypothesis, we found that all three copies of the putative genes involved in the biosynthesis of CNglc are expressed during adulthood (Figure 4) and mature adults contain twice as much CNglc as virgin adults (Figure 3a).

Cardoso and Gilbert, (2013) did not observe an increased cyanide emission from imagines of *H. charithonia, H. ethila* and *H. hecale* fed with an amino acid-rich diet in comparison with those fed only a sugar-solution in the first 2 weeks after eclosion. However, Nahrstedt and Davis, (1985) found that adults of *H. melpomene* continue to synthesize CNglcs even on a diet lacking amino acids, but their production tend to decline from 14 to 28 days and mortality increases. Perhaps a diet supplied with amino acids, such as pollen, is more important for the CNglc biosynthesis in the late period of the adult stage in *Heliconius*, when it is also crucial for their survivorship. They might initially be able to rely on the CNglcs obtained at the larval stage and maintain biosynthesis of linamarin and lotaustralin with amino acids from protein degradation.

Even though eggs neither biosynthesise nor sequester CNglcs, they had a similar total concentration of linamarin and lotaustralin as compared to larvae in late instars, parental pupae and newly emerged imagines (Figure 3a). Nahrstedt and Davis (1983) also found that the concentration of cyanide emitted by eggs of the heliconiine species *A. vanillae* and *D. iulia* resembled the concentration found imagines. These results show that heliconiines considerably invest in the cyanogenic content of its progeny by transferring CNglcs to the eggs.

Although concentration of CNglcs (linamarin and lotaustralin) decreased during the transition from eggs to early larvae (Figure 3a), the total amount of CNglcs was the same in both stages (3.18 ± 0.57 µg in eggs; 2.97 ± 0.56 µg in early larvae), indicating that this reduction in concentration was possibly a dilution effect. An intense expression of *CYP405A4* was observed in larvae at the early instars (Figure 4a), but it was not followed by the expression of *CYP332A1*, the gene putatively responsible for the conversion of the oxime to α-hydroxynitrile. *CYP405A4* and *CYP332A1* were both highly expressed in the gut transcriptome of fifth instar larvae (Figure 4b), which is in line with the observation that linamarin and lotaustralin levels are increased in the late as compared to early instar larvae, most likely due to de novo biosynthesis. qPCR and transcriptome results indicate that *CYP405A4* is the major active copy during larval stages and it is the most expressed in adults in comparison to *CYP405A5* and *CYP405A6.*

### 4.4. Linamarin and lotaustralin are used as nuptial gifts

In the present study linamarin and lotaustralin, but not dihydrogynocardin, were found in the spermatophore of *H. melpomene*, indicating that these CNglcs function as nuptial gifts (Figure 3b). Even though male and female adults contained dihydrogynocardin (Figure 3a), this compound was not present in their spermatophores and testes (Figure 3b), and it was not transferred to the eggs (Figure 3a). It is possible that testes and/or accessory glands produce the linamarin and lotaustralin transferred to the spermatophores or that only these CNglc can be transported to reproductive organs. Zagrobelny and collaborators (2014) demonstrated that radiolabelled linamarin and lotaustralin sequestered by *Z. filipendulae* larvae were distributed to all tissues during pupation and also transferred to the females as nuptial gifts during copula. Small amounts of radiolabelled linamarin and lotaustralin were even retrieved from some eggs produced by these females. However, sequestration of linamarin and lotaustralin is rare in *heliconiine*, and in general, they rely on obtaining aliphatic CNglcs via biosynthesis and not sequestration (Castro et al., 2019).

Although dihydrogynocardin was neither used as nuptial gifts nor transferred to eggs in our experiment with *H. melpomene*, it was consumed during the adult life time. How and why the butterflies catabolize the dihydrogynocardin obtained from larval sequestration are unanswered questions and further investigations are necessary. Perhaps they degrade dihydrogynocardin to release volatile cyanide and ketones/aldehydes during calling and courtship behaviour, as seen in *Z. filipendulae* (Zagrobelny et al., 2015). The cyanide emitted by these moths might allow them to estimate the CNglc content of their peers, as females of *Z. filipendulae* will only copulate with males having a high CNglc content. Rejected males having a low CNglc content were only accepted for mating after injection with linamarin or lotaustralin (Zagrobelny et al., 2015). Many *Heliconius species* use chemical signals for mating choice, as sex pheromones (Mann et al., 2017; Darragh et al., 2017) and antiaphrodisiacs (Gilbert, 1976; Schulz et al., 2007), but it is still unknown if the degradation products of CNglcs are part of these bouquets.

CNglcs seem to play an important role in reproduction in lepidopteran species that produce and/or sequester these compounds. Experiments regarding the CNglc composition of spermatophores of other *Heliconius* species and also from several successive matings would improve our understanding of the sexual selection and evolution of mating strategies in the genus.

### 4.5. CNglcs in Heliconius – functional diversification and interplay of multiple pathways

In this study, we demonstrated that *H. melpomene* larvae can sequester aliphatic, aromatic and cyclopentenyl CNglcs from *Passiflora.* Nevertheless, some types of CNglcs are more efficiently taken up than others suggesting that the larval ability to sequester different CNglc structures may be caused by transporters with low substrate specificity.

*H. melpomene* is most cyanogenic as mature adults due to intense biosynthesis of linamarin and lotaustralin in the adult stage. Predation by birds and lizards could have favoured CNglc accumulation during the adult stage, which is supported by the aposematic coloration and Müllerian mimicry observed in the *Heliconius* genus, and the long half-life of *H. melpomene*. However, since they use these compounds as nuptial gifts, sexual selection could have led these butterflies to increase their CNglc content as adults as well, especially in species of the non-pupal mating clade.

Cyclopentenyl CNglcs obtained during the larval stages were turned-over by mature adults, and since these compounds were not transferred to mating partners and neither to their progeny, we hypothesize that they catabolize these CNglcs to release sugar and nitrogen (turn-over) and/or use their volatile degradation products as essential mating cues as seen for aliphatic CNglcs in *Z. filipendulae* (Zagrobelny et al., 2015).

Although there is no doubt that CNglcs play a very important role in the biology of *Heliconius* species, the enzymes involved in the biosynthesis, sequestration, catabolism and detoxification of these compounds remain to be identified in these butterflies. The discovery and exploration of these pathways in the future will greatly expand our understanding of the roles and regulation of CNglcs in insects in general and in butterflies in particular.

## Acknowledgement

We thank the *Independent Research Fund Denmark | Natural Sciences* (DFF – 1323-00088) and the *Brazilian National Council for Scientific Development* (CNPq – proc. 306985/2013-6) for the financial support. EC acknowledge the *Science Without Borders* (CAPES) for her PhD scholarship and the Marie Curie Actions for her postdoctoral fellowship (Acronym: Cyanide Evolution). AO was funded by the *Cambridge Trust*, the *Cambridge Natural Environment Research Council Doctoral Training Partnership* and *St. John’s College Cambridge*. EC and AO would like to thank Professor Chris Jiggins for the additional financial support through the European Research Council grant number 339873 (Acronym: SpeciationGenetics). We thank Dr. Stefan Pentzold and Dr. Thiago Barbosa Cahu for the assistance in the larval frass and gut pH analyses, respectively. We are also grateful for all suggestions made by the Plant Biochemistry section (University of Copenhagen – PLEN) and the *Heliconius* Research Community to this project.

## Graphic Abstract

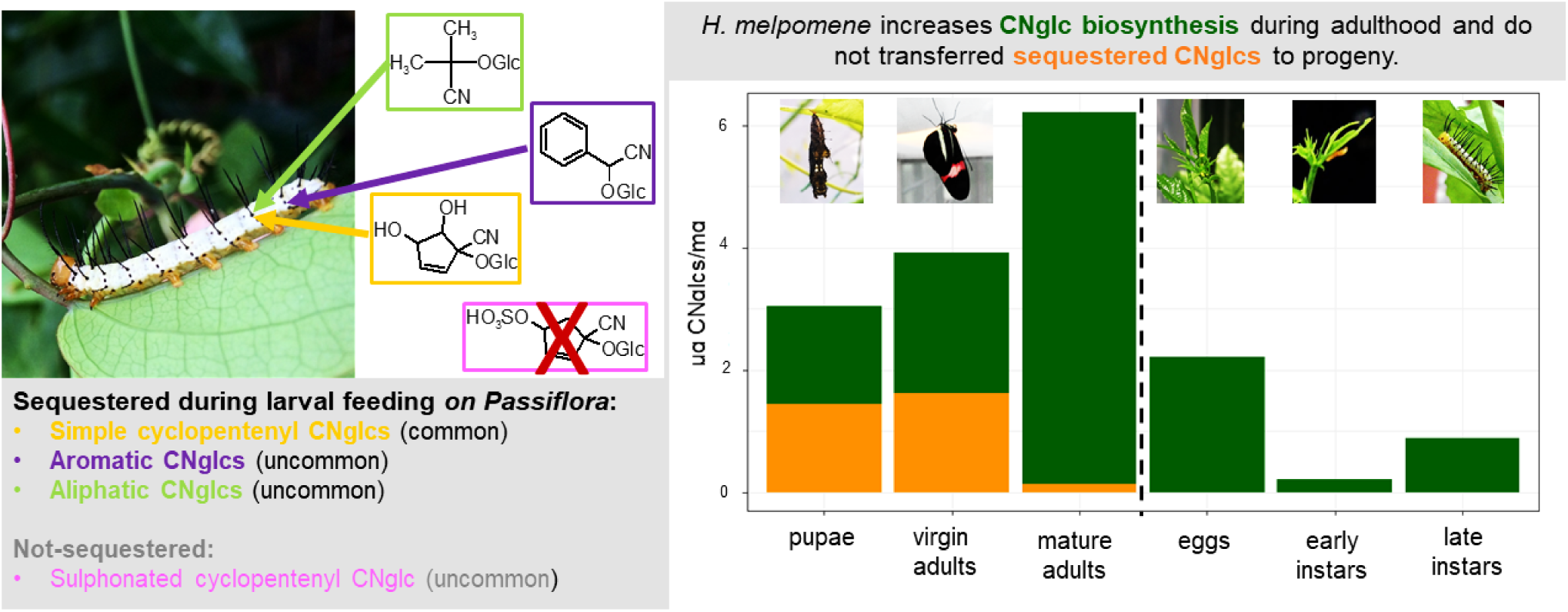

### Highlights

- *Heliconius melpomene* larvae can sequester not only cyclopentenyl, but also aliphatic and aromatic cyanogenic glucosides from *Passiflora* plants.
- Addition of sulphate groups to cyclopentenyl cyanogenic glucosides prevents sequestration – possibly a recent *Passiflora* adaptation to circumvent heliconiine herbivory.
- Biosynthesis of aliphatic cyanogenic glucoside is intensified in *H. melpomene* adults, when all three *CYP405A* genes were expressed.
- Sequestered cyclopentenyl cyanogenic glucosides are catabolized during the adult stage in *H. melpomene*.
- In contrast to linamarin and lotaustralin, sequestered cyclopentenyl cyanogenic glucosides are neither used as nuptial gifts nor transferred to the offspring.

## Supplementary Tables

**Table S1.**
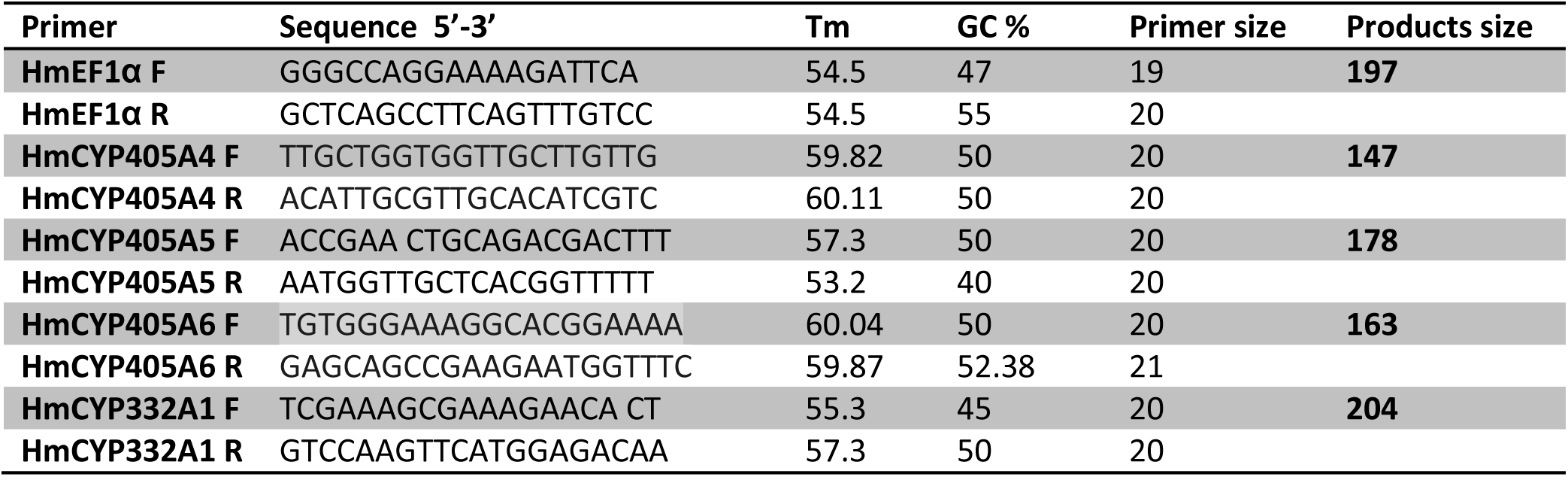
Primer sets utilized for qRT-PCR

**Table S2.** qPCR program. Step 2-4 were carried out in 40 cycles.

| Steps | Temperature | Time |
| --- | --- | --- |
| 1 | 95 °C | 7 min |
| 2 | 95 °C | 10 sec |
| 3 | 60 °C | 30 sec |
| 4 | 60 °C | 1 min |
| 5 | 60 °C to 98 °C<br>(0.5 °C increment per cycle) | 5 sec |

## Bibliography

Alfonso, J., Ortego, F., Sanchez-Monge, R., Garcia-Casado, G., Pujol, M., Castañera, P., Salcedo, G., 1997. Wheat and Barley Inhibitors Active Towards -Amylase and Trypsin-like Activities from Spodoptera frugiperda. J. Chem. Ecol. 23, 1729–1741. https://doi.org/10.1023/B:JOEC.0000006447.85514.8d

Alonso Amelot, M.E., Luis Ávila Núñez, J., Duarte, L., Oliveros-Bastidas, A., 2006. Hydrogen cyanide release during feeding of generalist and specialist lepidopteran larvae on a cyanogenic plant, Passiflora capsularis. Physiol. Entomol. 31, 307–315. https://doi.org/10.1111/j.1365-3032.2006.00528.x

Berenbaum, M., 1980. Adaptive Significance of Midgut pH in Larval Lepidoptera. Am. Nat. 115, 138–146. https://doi.org/Doi10.1086/283551

Boggs, C.L.., 1981. Selection Pressures Affecting Male Nutrient Investment at Mating in Heliconiine Butterflies. Evolution (N. Y). 35, 931–940.

Brown, K.S., 1981. The Biology of Heliconius and Related Genera. Annu. Rev. Entomol. 26, 427–457. https://doi.org/10.1146/annurev.en.26.010181.002235

Cardoso, M.Z., 2019. The effect of insect cyanoglucosides on predation by domestic chicks. bioRxiv. https://doi.org/10.1101/662288

Cardoso, M.Z., Gilbert, L.E., 2013. Pollen feeding, resource allocation and the evolution of chemical defence in passion vine butterflies. J. Evol. Biol. 26, 1254–1260. https://doi.org/10.1111/jeb.12119

Cardoso, M.Z., Gilbert, L.E., 2007. A male gift to its partner? Cyanogenic glycosides in the spermatophore of longwing butterflies (Heliconius). Naturwissenschaften 94, 39–42. https://doi.org/10.1007/s00114-006-0154-6

Chauhan, R., Jones, R., Wilkinson, P., Pauchet, Y., Ffrench-Constant, R.H., 2013. Cytochrome P450-encoding genes from the Heliconius genome as candidates for cyanogenesis. Insect Mol. Biol. 22, 532–40. https://doi.org/10.1111/imb.12042

Christensen, J., Jaroszewski, J.W., 2001. Natural glycosides containing allopyranose from the passion fruit plant and circular dichroism of benzaldehyde cyanohydrin glycosides. Org. Lett. 3, 2193–2195. https://doi.org/10.1021/ol016044+

Darragh, K., Salazar, C., Gonzalez-Rojas, M.F., McMillan, W.O., Pardo-Diaz, C., Mann, F., Schulz, S., Morrison, C.R., Darragh, K., Merrill, R.M., Vanjari, S., Jiggins, C.D., 2017. Male sex pheromone components in Heliconius butterflies released by the androconia affect female choice. PeerJ 5, e3953. https://doi.org/10.7717/peerj.3953

Davey, J.W., Chouteau, M., Barker, S.L., Maroja, L., Baxter, S.W., Simpson, F., Merrill, R.M., Joron, M., Mallet, J., Dasmahapatra, K.K., Jiggins, C.D., 2016. Major Improvements to the Heliconius melpomene Genome Assembly Used to Confirm 10 Chromosome Fusion Events in 6 Million Years of Butterfly Evolution. G3&#58; Genes|Genomes|Genetics. https://doi.org/10.1534/g3.115.023655

Davis, R.H., Nahrstedt, A., 1987. BIOSYNTHESIS OF CYANOGENIC GLUCOSIDES. Insect Biochem. 17, 689–693.

de Castro, É.C.P., Zagrobelny, M., Cardoso, M.Z., Bak, S., 2018. The arms race between heliconiine butterflies and Passiflora plants – new insights on an ancient subject. Biol. Rev. 93, 555–573. https://doi.org/10.1111/brv.12357

de Castro, É.C.P., Zagrobelny, M., Zurano, J.P., Cardoso, M.Z., Feyereisen, R., Bak, S., 2019. Sequestration and biosynthesis of cyanogenic glucosides in passion vine butterflies and consequences for the diversification of their host plants. Ecol. Evol. 9, 5079–5093. https://doi.org/10.1002/ece3.5062

Dobin, A., Davis, C.A., Schlesinger, F., Drenkow, J., Zaleski, C., Jha, S., Batut, P., Chaisson, M., Gingeras, T.R., 2013. STAR: ultrafast universal RNA-seq aligner. Bioinformatics 29, 15–21.

Ehrlich, P.R. &, Raven, R.J., 1964. Butterflies and Plants?: A Study in Coevolution Author. Soc. Study Evol. 18, 586–608.

Eisner, T., Eisner, M., Rossini, C., Iyengar, V.K., Roach, B.L., Benedikt, E., Meinwald, J., 2000. Chemical defense against predation in an insect egg. Proc. Natl. Acad. Sci. 97, 1634–1639. https://doi.org/10.1073/pnas.030532797

Engler-Chaouat, H.S., Gilbert, L.E., 2007. De novo synthesis vs. sequestration: negatively correlated metabolic traits and the evolution of host plant specialization in cyanogenic butterflies. J. Chem. Ecol. 33, 25–42. https://doi.org/10.1007/s10886-006-9207-8

Engler, H.S., Spencer, K.C., Gilbert, L.E., 2000. Preventing cyanide release from leaves. Nature 406, 144–5. https://doi.org/10.1038/35018159

Fitzgerald, T.D., 2008. Larvae of the fall webworm, Hyphantria cunea, inhibit cyanogenesis in Prunus serotina. J. Exp. Biol. 211, 671–677. https://doi.org/10.1242/jeb.013664

Fürstenberg-Hägg, J., Zagrobelny, M., Olsen, C.E., Jørgensen, K., Møller, B.L., Bak, S., 2014. Transcriptional regulation of de novo biosynthesis of cyanogenic glucosides throughout the life-cycle of the burnet moth Zygaena filipendulae (Lepidoptera). Insect Biochem. Mol. Biol. 49, 80–9. https://doi.org/10.1016/j.ibmb.2014.04.001

Gilbert, L.E., 1991. Biodiversity of a Central American Heliconius community: pattern, process, and problems, in: Price, P., Lewinsohn, T., Fernandes, T., Benson, W. (Eds.), Plant-Animal Interactions: Evolutionary Ecology in Tropical and Temperate Regions. John Wiley & Sons, New York, pp. 403–427.

Gilbert, L.E., 1976. Postmating female odor in Heliconius butterflies: A male-contributed antiaphrodisiac? Science (80-.). 193, 419–420. https://doi.org/10.1126/science.935877

Gilbert, L.E., 1972. Pollen feeding and reproductive biology of heliconius butterflies. Proc. Natl. Acad. Sci. U. S. A. 69, 1403–7.

Jaroszewski, J.W., Olafsdottir, E.S., Wellendorph, P., Christensen, J., Franzyk, H., Somanadhan, B., Budnik, B. a, Jørgensen, L.B., Clausen, V., 2002. Cyanohydrin glycosides of Passiflora: distribution pattern, a saturated cyclopentane derivative from P. guatemalensis, and formation of pseudocyanogenic alpha-hydroxyamides as isolation artefacts. Phytochemistry 59, 501–11.

Jensen, N.B., Zagrobelny, M., Hjernø, K., Olsen, C.E., Houghton-Larsen, J., Borch, J., Møller, B.L., Bak, S., 2011. Convergent evolution in biosynthesis of cyanogenic defence compounds in plants and insects. Nat. Commun. 2, 273. https://doi.org/10.1038/ncomms1271

Krueger, F., 2015. Trim galore. A wrapper tool around Cutadapt FastQC to consistently apply Qual. Adapt. trimming to FastQ files.

Liao, Y., Smyth, G.K., Shi, W., 2013. featureCounts: an efficient general purpose program for assigning sequence reads to genomic features. Bioinformatics 30, 923–930.

Mann, F., Vanjari, S., Rosser, N., Mann, S., Dasmahapatra, K.K., Corbin, C., Linares, M., Pardo-Diaz, C., Salazar, C., Jiggins, C., Schulz, S., 2017. The Scent Chemistry of Heliconius Wing Androconia. J. Chem. Ecol. 43, 843–857. https://doi.org/10.1007/s10886-017-0867-3

Mason, P.A., Singer, M.S., 2015. Defensive mixology: Combining acquired chemicals towards defence. Funct. Ecol. https://doi.org/10.1111/1365-2435.12380

Møller, B.L., 2010. Functional diversifications of cyanogenic glucosides. Curr. Opin. Plant Biol. 13, 338–47. https://doi.org/10.1016/j.pbi.2010.01.009

Nabity, P.D., Orpet, R., Miresmailli, S., Berenbaum, M.R., DeLucia, E.H., 2012. Silica and Nitrogen Modulate Physical Defense Against Chewing Insect Herbivores in Bioenergy Crops <I>Miscanthus</I> × <I>giganteus</I> and <I>Panicum virgatum</I> (Poaceae). J. Econ. Entomol. 105, 878–883. https://doi.org/10.1603/EC11424

Nahrstedt, a., Davis, R.H., 1985. Biosynthesis and quantitative relationships of the cyanogenic glucosides, linamarin and lotaustralin, in genera of the Heliconiini (Insecta: Lepidoptera). Comp. Biochem. Physiol. Part B Comp. Biochem. 82, 745–749. https://doi.org/10.1016/0305-0491(85)90519-X

Nahrstedt, A., Davis, R.H., 1983. Occurrence, variation a n d biosynthesis of the cyanogenic glucosides linamarin a n d lotaustralin in species of the. Comp. Biochem. Physiol. 75, 65–73. http://dx.doi.org/10.1016/0305-0491(83)90041-X

Nishida, R., 2002. Swquestration of Defensive Substances from Plants by Lepidoptera. Annu. Rev. Entomol. 47, 57–92.

Pentzold, S., Zagrobelny, M., Roelsgaard, P.S., Møller, B.L., Bak, S., 2014a. The multiple strategies of an insect herbivore to overcome plant cyanogenic glucoside defence. PLoS One 9, e91337. https://doi.org/10.1371/journal.pone.0091337

Pentzold, S., Zagrobelny, M., Rook, F., Bak, S., 2014b. How insects overcome two-component plant chemical defence: Plant b-glucosidases as the main target for herbivore adaptation. Biol. Rev. 89, 531–551. https://doi.org/10.1111/brv.12066

Reed, R.D., Chen, R., Halder, G., Nijhout, H.F., McMillan, W.O., Papa, R., Martin, A., Hines, H.M., Counterman, B.A., Pardo-Diaz, C., Jiggins, C.D., Chamberlain, N.L., Kronforst, M.R., 2011. optix Drives the Repeated Convergent Evolution of Butterfly Wing Pattern Mimicry. Science (80-.). 333, 1137–1141. https://doi.org/10.1126/science.1208227

Robinson, M.D., McCarthy, D.J., Smyth, G.K., 2010. edgeR: a Bioconductor package for differential expression analysis of digital gene expression data. Bioinformatics 26, 139–140.

Schappert, P.J., Shore, J.S., 1999. Effects of cyanogenesis polymorphism in Turnera ulmifolia on Euptoieta hegesia and potential Anolis predators. J. Chem. Ecol. 25, 1455–1479. https://doi.org/Doi10.1023/A:1020995329 980

Schulz, S., Boppré, M., Yildizhan, S., Estrada, C., Gilbert, L.E., 2007. An Antiaphrodisiac in Heliconius melpomene Butterflies. J. Chem. Ecol. 34, 82–93. https://doi.org/10.1007/s10886-007-9393-z

Sheppard, P.M., Turner2-4, J.R.G., Brown, K.S., Benson, W.W., Singer, M.C., 1985. GENETICS AND THE EVOLUTION OF MUELLERIAN MIMICRY IN HELICONIUS BUTTERFLIES, Trans. R. Lond. B.

Spencer, K.C., 1988. Chemical Mediation of Coevolution in the Passiflora–Heliconius Interaction, in: Chemical Mediation of Coevolution. Elsevier, pp. 167–240. https://doi.org/10.1016/B978-0-12-656855-4.50011-5

Spencer, K.C., Seigler, D.S., Nahrstedt, A., 1986. Linamarin, lotaustralin, linustatin and neolinustatin from Passiflora species. Phytochemistry 25, 645–647. https://doi.org/10.1016/0031-9422(86)88016-5

Walters, J.R., Hardcastle, T.J., Jiggins, C.D., 2015. Sex chromosome dosage compensation in heliconius butterflies: Global yet still incomplete? Genome Biol. Evol. https://doi.org/10.1093/gbe/evv156

Wosch, L., Imig, D.C., Cervi, A.C., Moura, B.B., Budel, J.M., de Moraes Santos, C.A., 2015. Comparative study of Passiflora taxa leaves: I. A morpho-anatomic profile. Rev. Bras. Farmacogn. 25, 328–343. https://doi.org/10.1016/j.bjp.2015.06.004

Yu, Q.Y., Fang, S.M., Zhang, Z., Jiggins, C.D., 2016. The transcriptome response of Heliconius melpomene larvae to a novel host plant. Mol. Ecol. 25, 4850–4865. https://doi.org/10.1111/mec.13826

Zagrobelny, M., Bak, S., Ekstrøm, C.T., Olsen, C.E., Møller, B.L., 2007a. The cyanogenic glucoside composition of Zygaena filipendulae (Lepidoptera: Zygaenidae) as effected by feeding on wild-type and transgenic lotus populations with variable cyanogenic glucoside profiles. Insect Biochem. Mol. Biol. 37, 10–8. https://doi.org/10.1016/j.ibmb.2006.09.008

Zagrobelny, M., Bak, S., Møller, B.L., 2008. Cyanogenesis in plants and arthropods. Phytochemistry 69, 1457–68. https://doi.org/10.1016/j.phytochem.2008.02.019

Zagrobelny, M., Bak, S., Olsen, C.E., Møller, B.L., 2007b. Intimate roles for cyanogenic glucosides in the life cycle of Zygaena filipendulae (Lepidoptera, Zygaenidae). Insect Biochem. Mol. Biol. 37, 1189–97. https://doi.org/10.1016/j.ibmb.2007.07.008

Zagrobelny, M., Jensen, M.K., Vogel, H., Feyereisen, R., Bak, S., 2018. Evolution of the Biosynthetic Pathway for Cyanogenic Glucosides in Lepidoptera. J. Mol. Evol. 1. https://doi.org/10.1007/s00239-018-9854-8

Zagrobelny, M., Olsen, C.E., Pentzold, S., Fürstenberg-Hägg, J., Jørgensen, K., Bak, S., Møller, B.L., Motawia, M.S., 2014. Sequestration, tissue distribution and developmental transmission ofcyanogenic glucosides in a specialist insect herbivore. Insect Biochem. Mol. Biol. 44, 44–53. https://doi.org/10.1016/j.ibmb.2013.11.003

Zagrobelny, M., Simonsen, H.T., Olsen, C.E., Bak, S., Møller, B.L., 2015. Volatiles from the burnet moth Zygaena filipendulae (Lepidoptera) and associated flowers, and their involvement in mating communication. Physiol. Entomol. 40, 284–295. https://doi.org/10.1111/phen.12113

